# In-field whole plant maize architecture characterized by Latent Space Phenotyping

**DOI:** 10.1101/763342

**Authors:** Joseph L. Gage, Elliot Richards, Nicholas Lepak, Nicholas Kaczmar, Chinmay Soman, Girish Chowdhary, Michael A. Gore, Edward S. Buckler

## Abstract

Collecting useful, interpretable, and biologically relevant phenotypes in a resource-efficient manner is a bottleneck to plant breeding, genetic mapping, and genomic prediction. Autonomous and affordable sub-canopy rovers are an efficient and scalable way to generate sensor-based datasets of in-field crop plants. Rovers equipped with light detection and ranging (LiDar) can produce three-dimensional reconstructions of entire hybrid maize fields. In this study, we collected 2,103 LiDar scans of hybrid maize field plots and extracted phenotypic data from them by Latent Space Phenotyping (LSP). We performed LSP by two methods, principal component analysis (PCA) and a convolutional autoencoder, to extract meaningful, quantitative Latent Space Phenotypes (LSPs) describing whole-plant architecture and biomass distribution. The LSPs had heritabilities of up to 0.44, similar to some manually measured traits, indicating they can be selected on or genetically mapped. Manually measured traits can be successfully predicted by using LSPs as explanatory variables in partial least squares regression, indicating the LSPs contain biologically relevant information about plant architecture. These techniques can be used to assess crop architecture at a reduced cost and in an automated fashion for breeding, research, or extension purposes, as well as to create or inform crop growth models.

---

The cost to genotype a population of plants has become increasingly affordable since the advent of next-generation sequencing, leaving the acquisition of high quality phenotypic data as a limiting step in conducting genetic mapping studies and training genomic prediction models. Sensors and imaging devices with the ability to capture terabytes of data, rather than providing a solution to the phenotyping bottleneck, can compound the problem by producing hundreds or thousands of non-independent phenotypes for downstream analysis or, at worst, producing massive data sets that are never used to their full potential. Novel data acquisition methods require custom computational methods for extracting interpretable, useful, and biologically relevant traits from this deluge of digital data.

The massive quantity of data produced by image- and sensor-based phenotyping methods has created new challenges regarding data management and analysis (Omasa et al., 2007; Houle et al., 2010; Minervini et al., 2015). Many analysis methods for novel phenotyping methods are lab-oriented, site specific, or cannot be directly applied to crops in situ (Araus et al., 2018). To be useful, phenotypes derived from new imaging and sensor methods need to be applicable in breeding programs or useful for gaining biological insights (Cobb et al., 2013). Many recently developed high throughput phenotyping platforms are immobile (Virlet et al., 2017; Bai et al., 2019) or mounted on tractors (Busemeyer et al., 2013; Andrade-Sanchez et al., 2014; Salas Fernandez et al., 2017; Sun et al., 2018). Such solutions are viewed with some skepticism by plant breeders, who prioritize flexible, affordable approaches to survey large populations of germplasm in numerous growing environments (Araus et al., 2018). An ideal phenotyping system should have high mobility/flexibility, high sensor capacity, be easily scaled up, and be cheap (Figure 1). Unmanned aerial vehicles (UAVs) and ground rovers meet most of those needs. However, UAVs can be difficult to fly in poor weather (Yang et al., 2017) and are limited in their ability to characterize plant architecture below and within the canopy, especially late in the growing season (Sun et al., 2018). Equipped with RGB cameras and LiDar, rovers can be inexpensive relative to existing automated phenotyping systems, making them scalable and flexible enough to characterize large populations in many locations. Because they are small enough to fit between conventionally planted rows of maize, they can be used to characterize plant architecture from a perspective that is not accessible by most existing phenotyping systems (Mueller-Sim et al., 2017; Higuti et al., 2018; Kayacan et al., 2018; Stager et al., 2019). LiDar sensors enable 3-dimensional reconstruction of field plots, which can be used to characterize plant architecture and biomass distribution.

**Figure 1:**
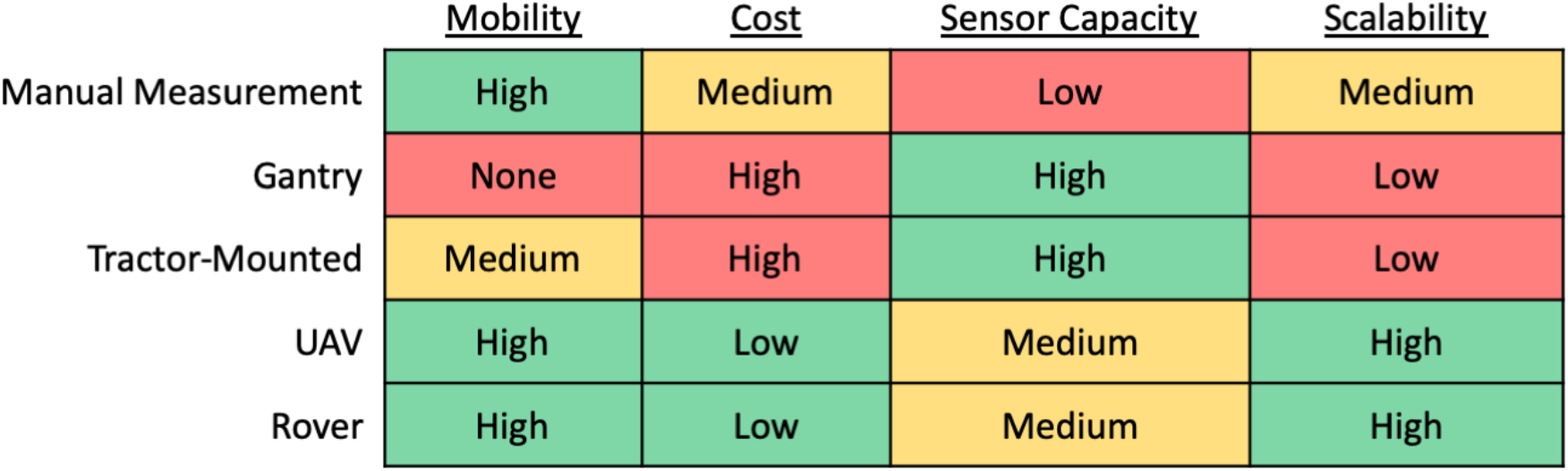
Comparison of phenotyping methods. To be effectively implemented in breeding programs or large-scale biological studies, phenotyping technologies need to be mobile, cheap, and scalable while enabling collection of relevant data by onboard (or in-hand) sensors. Rovers and UAVs meet most of those requirements. For within- and below-canopy phenotyping, rovers have an advantage over UAVs, which can only characterize trials from above.

Three-dimensional reconstructions of in-field crop plots, though information-rich, also present challenges for analysis. Methods for analyzing image or sensor data are often created for a specific purpose or study, and can be difficult for external groups to apply to newly acquired datasets. These issues can occur due to threshold or parameter specification, image settings or modality, or ad-hoc solutions that do not translate well to new datasets (Ubbens et al., 2019). Latent Space Phenotyping (LSP), described by Ubbens et al. (2019), leverages the flexibility of machine learning to address some of the inherent difficulties of analyzing and extracting phenotypes from large, complex, sensor- or image-based datasets. Rather than develop methods to measure human-defined traits such as plant height or leaf number, Ubbens and colleagues (2019) describe LSP as a way to generate phenotypes that are an “abstract learned concept of the response to treatment, inferred automatically from the image data using deep learning” (Ubbens et al., 2019). Described another way, LSP allows a computer to process images or sensor-based data in such a way as to produce novel traits that distinguish individuals based on treatments. As an example, Ubbens and colleagues (2019) used LSP on RGB images of well-watered and water-limited *Setaria viridis* plants from a biparental population to create and evaluate computer-generated traits that distinguish the two treatments based on characteristics learned from the images. They used those traits for QTL mapping and were able to reproduce discovery of a QTL related to water use (Ubbens et al., 2019). In this study, we expand their definition to also include principal component analysis (PCA), which provides a way to create latent phenotypes without building machine learning models. We evaluate both PCA and a machine learning model called an autoencoder, both of which generate Latent Space Phenotypes (LSPs) that, rather than distinguishing between treatments, distinguish between different varieties of hybrid maize.

PCA is a method of dimensionality reduction that creates latent variables from linear combinations of the original data. These latent variables, or principal components (PCs), are ordered, orthogonal to each other, and greedy such that early PCs capture more variability than later PCs. By using PCA, an 8,192-dimensional image (64 pixels by 128 pixels), where many of the dimensions are correlated with each other (e.g., adjacent pixels), can be compressed to a reduced-dimensional representation described by a chosen number of independent PCs. As such, PCA serves as a way to both reduce dimensionality and impose independence between variables.

Autoencoders are a type of unsupervised neural network that can be trained to learn a reduced representation of their input (Rumelhart et al., 1986; Baldi, 2012). Conceptually similar to various image compression methods, this reduced representation can be used to reconstruct, with some loss of information, the original input. The difference between the original input and the reconstructed version can be used as a heuristic to train the network. Depending on their architecture, autoencoders can produce results similar to PCA. By adding convolutional layers and non-linear activations to the autoencoder, however, it can capture different relationships between image elements than PCA, which is based entirely on linear combinations of pixel values.

One of the main challenges in producing LSPs from LiDar data is to extract signal while ignoring noise. Data from each plot consists of hundreds of thousands of data points, and the data are inherently noisy due to rover position and velocity during data acquisition, plant movement, and sensor error. Useful LSPs will contain information about plant architecture and plot-level biomass distribution regardless of rover position, wind speed and direction, or other factors that contribute noise. Because of this, the objective of the autoencoder in this study is no longer to accurately recreate the input, but to recreate the elements of the input that are relevant to plant biomass while excluding or ‘ignoring’ signal that comes from unwanted sources of variability.

In this study, we evaluate both PCA and an autoencoder as methods to extract LSPs in an unsupervised manner from in-field LiDar scans of hybrid maize plots. We calculate heritability for the LSPs and compare them to phenotypes that are traditionally measured manually, showing that LSPs have similar heritability to traditionally evaluated phenotypes. LSPs are then used as explanatory variables to predict manually measured phenotypes, demonstrating that LSPs, though not directly interpretable, contain information about plant architecture and biomass distribution. By demonstrating heritability and relevance to plant architecture and biomass distribution, we show that LSPs are a useful tool with applications to plant breeding and biology.

## Materials & Methods

### Germplasm and Manual Phenotyping

The field experiments used in this study were the 2018 Genomes to Fields NYH2 and NYH3 locations. Genomes to Fields is a multi-year, multi-location coordinated experiment that was planted at 31 unique locations across the United States and Canada in 2018 (Gage et al., 2017; AlKhalifah et al., 2018; Lawrence-Dill et al., 2019). The NYH2 and NYH3 locations used for this study consisted of 1,600 two-row plots containing 890 unique maize hybrids planted in a modified randomized complete block design in Aurora, NY. Plots were 5.33m long with 0.76m space between rows and 1.07m alleys between ranges. The field was laid out as two experiments, NYH2 and NYH3, which were each 25 plots (50 rows) wide and 32 plots (ranges) long. Twenty hybrid checks and 84 photoperiod introgression lines crossed to LH123Ht, were planted in both NYH2 and NYH3. The remainder of each field consisted of three inbred biparental populations sharing a common parent (Mo44 x PHW65, PHN11 x PHW65, PHW65 x MoG; (Gage et al., 2018), crossed to either PHT69 (in NYH2) or LH195 (in NYH3). Manual measurements were recorded for days to silk, days to anthesis, plant height, ear height, stand count, root lodging, stalk lodging, grain moisture, plot weight, test weight, total leaf count, and leaf number up to the primary ear. All traits except the last two were evaluated according to the Genomes to Fields standard operating procedure (https://www.genomes2fields.org/resources/), with plant height and ear height measured on a single representative plant per plot. Leaf count traits were performed at maturity, by counting leaves on one representative plant per plot.

### Rover Design and Sensors

The TerraSentia rover is a compact, autonomous, and teachable robot designed by Earthsense, Inc. (https://www.earthsense.co) for high-throughput phenotyping beneath the canopy of agricultural crops. It is battery operated, and has an onboard Intel i7 computer with 500GB SSD and 16GB RAM as well as a separate autonomy computer. The rover is equipped with three RGB cameras and two planar Hokuyo UST-10LX LiDar that record continuously as it drives through a field (Kayacan et al., 2018). Furthermore, it has the capability to fit between rows in standard maize fields and can autonomously follow rows using an embedded LiDar (Higuti et al., 2018).

Data collection by the rovers took place after all lines had reached physiological maturity (flowering) but before harvest. All data were collected on September 4th, 7th, 12th, 19th or 20th, 2018. Rovers were driven along a single column of the field at a time, passing between the rows of two-row plots such that the data recorded on both sides of the rover at any given time corresponded to the same hybrid and the same plot. Data belonging to discrete plots within each column of the field were separated based on manually recorded time stamps of the video feed from the side-facing cameras. Incomplete passes of the field, and plots that were not driven through continuously, were manually removed from the dataset. The cleaned dataset (including border plots; numbers in parentheses exclude border plots) consisted of 2,103 (1,972) discrete plot-level records from 1,153 (1,083) unique plots (many plots were driven through and recorded more than once) representing 698 (697) unique hybrids. Border plots were included for the process of calculating LSPs, because they contain useful data that can be used during PCA and autoencoder training. They were not included in subsequent heritability calculations or predictions of manually measured phenotypes.

### Cleaning and Preprocessing LiDar Data

LiDar data for each plot were recorded every 25 milliseconds. Each time point was recorded as a vector of distances at 1,080 discrete angles around the LiDar unit. These can be processed as a 2D matrix with 1,080 rows and as many columns as time points within each plot, where each entry corresponds to a measured distance. The distances and angles at which the measurements were taken can also be converted from polar coordinates to cartesian coordinates for processing as a point cloud. Because the Hokuyo UST-10LX is a planar lidar, the data recorded at any given time point correspond to a two dimensional ‘slice’ within a plane perpendicular to the motion of the rover. A three dimensional representation of each plot can be created by arranging the slices along a third dimension corresponding to their timestamps as the rover drives through the plot (Figure 2).

**Figure 2:**
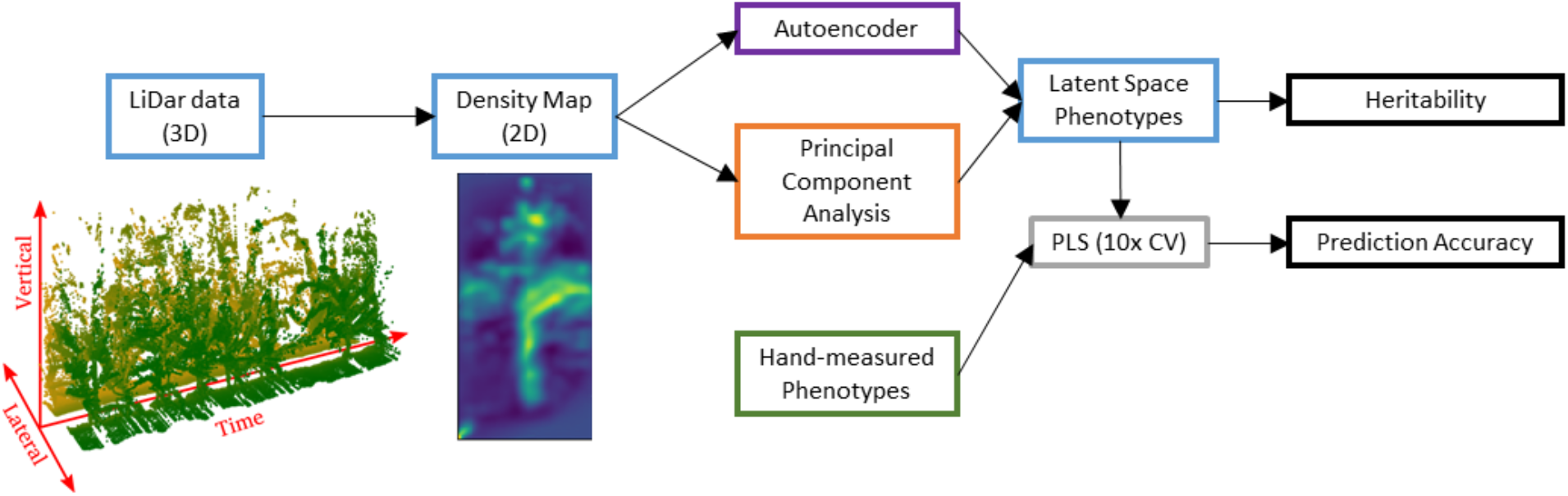
Schematic of this study’s workflow. 3-dimensional LiDar data are compressed into 2-dimensional density maps. Examples of a LiDar point cloud and density map can be seen below their respective labelled boxes. The density map images are subjected to dimensionality reduction by principal component analysis and an autoencoder, resulting in Latent Space Phenotypes (LSPs) that contain information about plant architecture and biomass distribution. Then the LSPs’ heritability were calculated, and they were used to predict manually measured phenotypes.

LiDar data cleaning and processing were performed in R (R Core Team, 2018) using the ‘ash’ (Scott et al., 2009), ‘EBImage’ (Pau et al., 2010), ‘mmand’ (Clayden, 2014), ‘slidaRtools’ (http://github.com/niknap/slidaRtools), and ‘tidyverse’ (Wickham, 2017) packages. Three dimensional point clouds were cleaned by first removing points outside of the plot of interest. Because LiDar measures distance to the nearest object in a straight line, many data points correspond to plants in adjacent plots. We removed any points outside the plot of interest by thresholding the absolute horizontal distance from the rover according to Otsu’s method (Otsu, 1979). Following this step, points belonging to the ground were identified by performing a morphological opening on the image with a 65×65 square kernel, which leaves only points belonging to the ground. The ground points were removed from the original image, and the matrix converted into three dimensional point cloud format. The point cloud was aggregated into voxels at a resolution of 5mm × 5mm within each time point. Voxels containing only a single LiDar data point and with no neighbors in three-dimensional space were assumed to be noise and were subsequently removed.

Cleaned, three-dimensional LiDar data for each distinct point cloud were then compressed into a two dimensional density map. First, the two rows of the plot were merged by reflecting one row of the plot around the vertical axis. This effectively lined all plants from both rows of the plot into a single row. Second, data points were binned along the length of the plot (represented by the time dimension; Figure 2) into 128×64 bins (vertical by lateral) and two dimensional density was calculated by average shifted histogram (Scott, 2015). This results in an image that resembles a single plant, representing the average distribution of all plant surfaces in the plot. Applying this compression to all plot-level point clouds resulted in 2,103 density maps that were subjected to dimensionality reduction to extract LSPs (Ubbens et al., 2019).

### Creating Latent Space Phenotypes

Two dimensionality reduction methods were applied to the plant surface density maps: PCA and an autoencoder. To format the data for PCA, each plant surface density map was vectorized, then row vectors from all observations were bound together into a matrix. The density values were log-transformed, then centered and scaled column-wise (pixel-wise), before performing PCA. The principal component scores for each plant surface density map were used as PCA-based LSPs.

One of the primary goals of the autoencoder was to extract an encoded representation of each plot that represents plant architecture and biomass distribution, while ignoring variability that is attributable to differences in rover position, wind direction, or other factors that introduce noise. In some senses this resembles the goals of denoising autoencoders, which can be trained to remove noise from images, but in this case we did not know ahead of time which elements of the LiDar data constituted noise and which constituted signal, nor did we know the distribution of noise in order to simulate it. Because many plots were driven more than once by the rover, we were able to use those technical replications to identify features of a density map that are persistent between technical replications, while ignoring those that are variable due to noise. The autoencoder was constructed to take two plant surface density maps as input images, representing repeated data collections on the same plot. Both technical replication images are encoded separately, but their encodings are averaged before being decoded, in order to train the autoencoder to identify a reduced representation that can be used to reconstruct both images. Figure 3 has a schematic of the autoencoder model described below.

**Figure 3:**
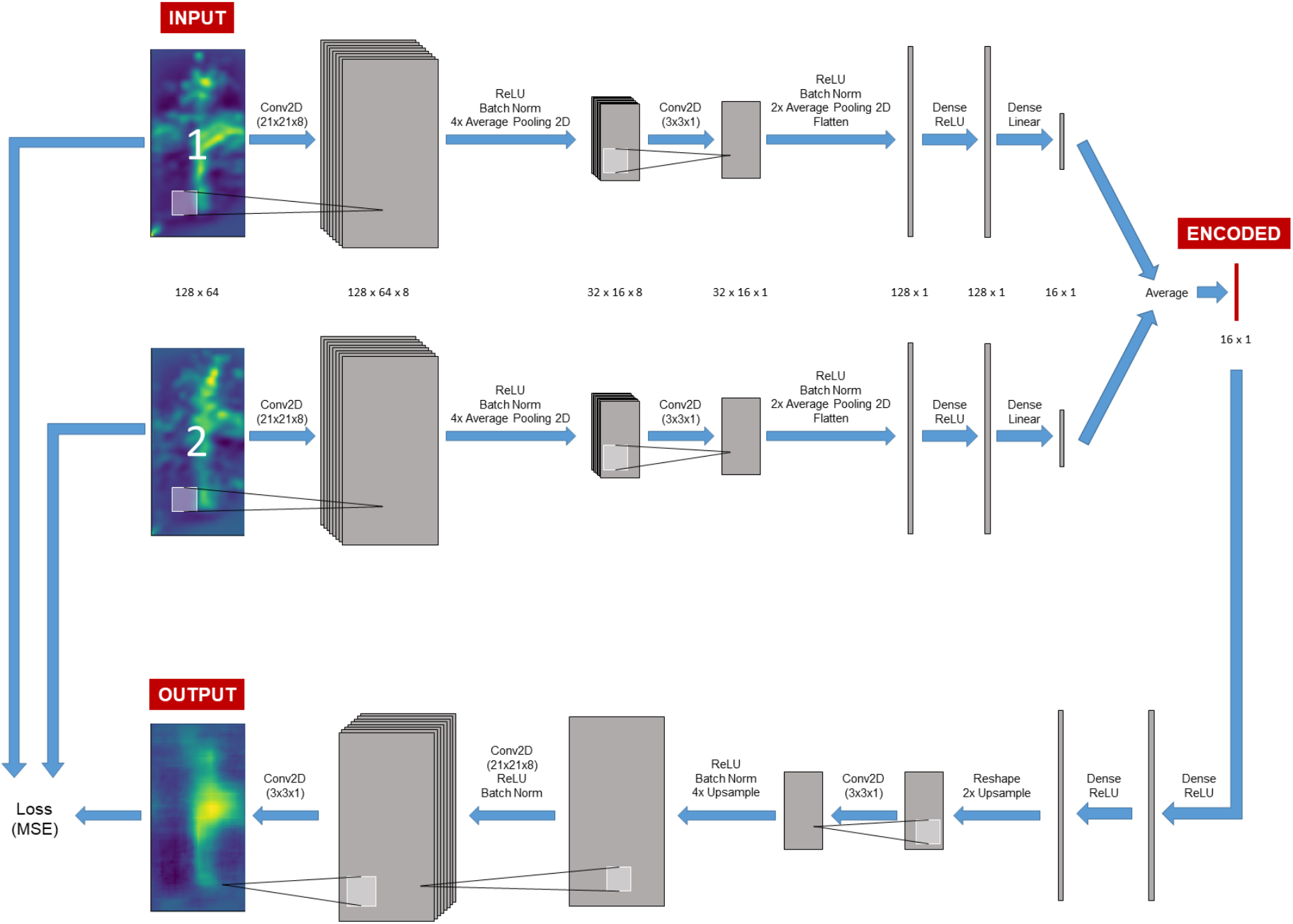
Diagram of autoencoder used for extracting Latent Space Phenotypes (LSPs). Many field plots were driven by the rover more than once, producing technical replications of data collection in those plots. Technical replications were used to train the autoencoder to identify signal that is persistent in both images, while ignoring noise caused by rover position, wind, or other stochastic influences. Both technical replications are put through identical encoding pipelines, where they are convolved twice and put through two dense layers to produce a 16-dimensional encoding. Before decoding, the encoded representations of the technical replicates are averaged. Decoding proceeds through two dense layers and three convolutional layers before producing the output image. Loss was calculated as the mean squared error between the two input images and the output. After model training, the input was changed from two different technical replication images to duplicates of a single density map in order to produce unique encodings for each uniquely acquired density map.

For the encoding step, 128×64 pixel plant surface density maps are put through a series of convolutional layers combined with average pooling to reduce the size of the image. The resulting 32×16 matrix is flattened and passed through a series of dense layers to produce the vector of 16 encoded values. To decode, the encoded values are passed through a series of dense layers before being reshaped into a 32×16 image, which is subsequently convolved and upsampled to produce the output. L2 kernel regularization was used in all convolutional layers except the last, as well as when creating the 16-value encoded vector. The lambda value for all L2 regularizations was set to 0.01.

Of the 2,103 plot-level observations recorded, 1,386 were technical replications of 693 unique plots. These technical replications were augmented by rotating and adding noise, then used to train the autoencoder (Figure 4). 20% of the training data, split by plots, were used as a validation set to assess overfitting. By forcing technical replications to have the same encoded values during training, the network learns to ignore noise or perturbations that occur during repeated sampling. The loss function during model training was the mean squared error between the decoded image and the original plant surface density image for each of the technical replicates. The autoencoder was trained with a batch size of 256 and trained for 200 epochs, at which point validation set loss was stabilized.

**Figure 4:**
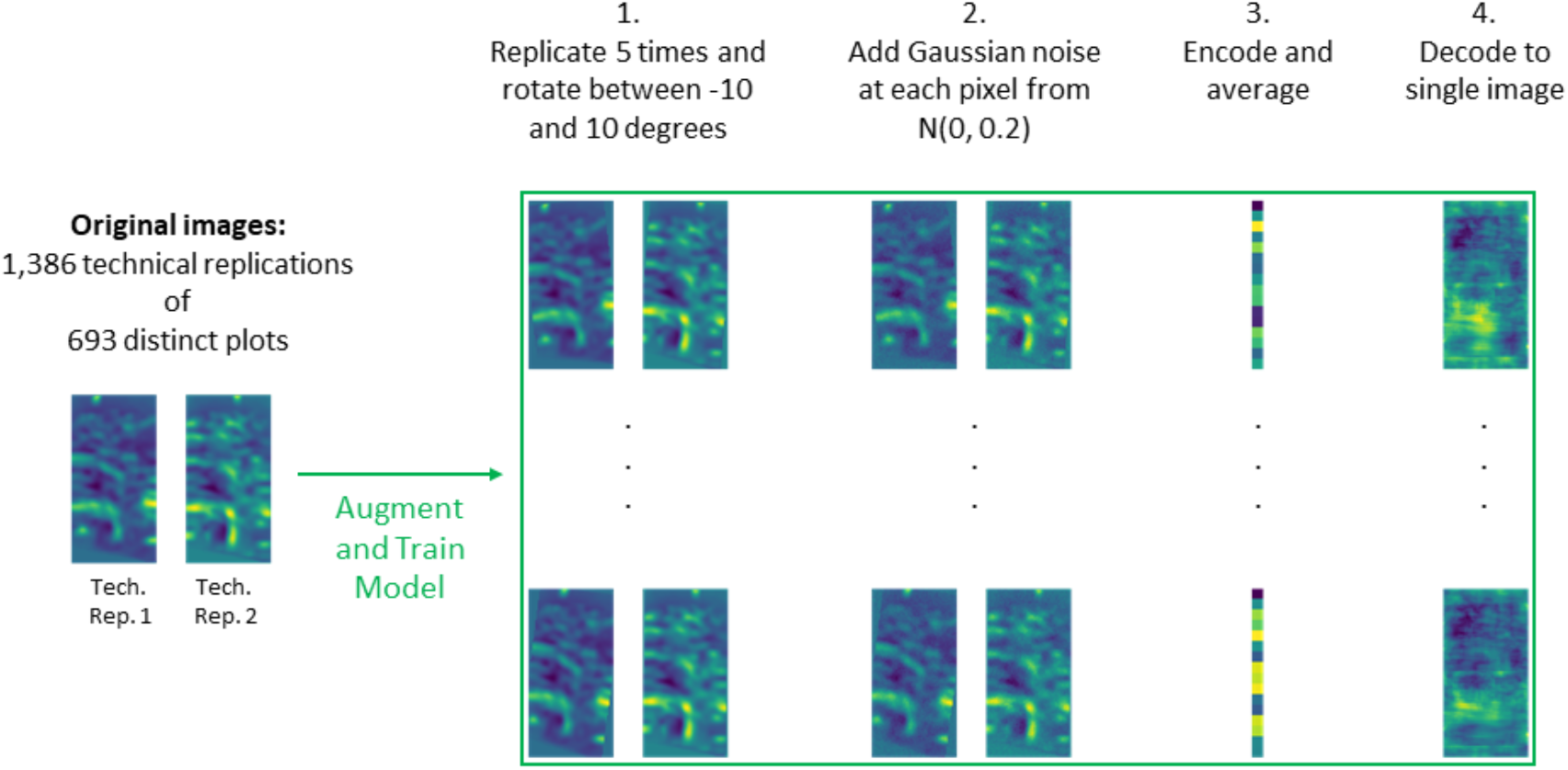
Image augmentation for model training. Distinct field plots that were recorded more than once represent technical replications. Because any two technical replications represent the same plot, they should have the same encoded values. Hence, for any given plot in the training set, the two technical replications were processed independently through the encoding portion of the network, but their encoded values were averaged before being decoded into a single image representing the plot. Density maps for training plots were replicated and augmented by random rotation and addition of noise.

After training, all original data was encoded. Unlike in training, replicated plots were encoded separately, allowing different encodings for each technical replication. Code for creating and training the model, as well as an HDF5 of the final model, can be found at https://bitbucket.org/bucklerlab/p_lidar_lsp.

### Evaluating Latent Space Phenotypes

LSPs were evaluated based on two metrics: their heritability and how well they predict manually measured phenotypes. If LSPs are heritable, it means there is genetic variation for these novel traits and they can be effectively selected on in an applied breeding program. If they can be used to predict manually measured phenotypes, it means the LSPs are not characterizing meaningless differences between individuals but instead contain information about plant architecture and other biologically important phenotypes.

Heritability was calculated by modelling manually measured phenotypes and LSPs as *y*_*ijk*_ = *μ* + *G*_*i*_ + *E*_*j*_ + *R*(*E*)_*kj*_ + *ɛ*_*ijk*_, where *y*_*ijk*_ is the manual phenotype or LSP of genotype *i* in experiment *j*, in block *k*; *μ* is the overall mean, *G*_*i*_ ~ *N*(0, *σ*^2^_*g*_*I*) is the random effect of the *i*th hybrid individual; *E*_*j*_ is the effect of the *j*th experiment (NYH2 or NYH3); *R*(*E*)_*kj*_ is the effect of block *k* nested in experiment *j*; and *ɛ*_*ijk*_ ~ *N*(0, *σ*^2^_*ɛ*_*I*) is a normally and independently distributed error term. Broad-sense heritability was calculated as 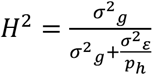, where *p*_*h*_ is the harmonic mean of the number of plots in which each hybrid was evaluated (Holland et al., 2003).

LSPs were used to predict manually measured phenotypes by partial least squares regression (PLSR). PLSR models were built using the *plsr()* function in the R (R Core Team, 2018) package ‘pls’ *(Wehrens and Mevik, 2007)* with 10-fold cross validation by hybrid and five latent variables. For each manually measured trait, separate predictions was performed using either the first 16 PCA LSPs or all autoencoder LSPs. Accuracy of the PLSR predictions was measured as the Pearson’s correlation between predicted and manually measured phenotypes.

We also used PLSR to see how well non-architectural traits can be predicted from manually measured architectural and biomass distribution traits alone. We used plant and ear height, total leaf number, ear leaf number, stand count, stalk lodging, and root lodging as explanatory variables to predict days to anthesis, days to silking, grain moisture, test weight, and plot weight using five latent variables and the same 10-fold cross validation scheme as above.

## Results and Discussion

Driving the rover through hybrid trials yielded 2,103 three-dimensional point cloud representations of 1,153 unique two-row plots containing 698 hybrid genotypes. Visual inspection of the point clouds reveals that they capture individual plant architecture well (see Figure 2 for an example point cloud). Previous to this study, there was some concern that because LiDar can only measure objects within “line of sight” of the sensor, the mid to upper canopy would be poorly represented due to occlusion. Rather, examination of individual plots revealed reconstruction of upper leaves and in some cases even tassels. By overlaying the two rows of a plot and calculating the density along the rows of the plot, we produced a two-dimensional “image” showing the distribution of plant surfaces for an average maize plant in that plot (see Figure 2 for an example density map). LSPs extracted from those plant surface distribution images by PCA or an autoencoder were evaluated for their heritability and whether they can be used to predict manually measured phenotypes.

### Latent Space Phenotype Heritability

We calculated the heritability of the LSPs in order to assess whether they are characterized by enough genetic variation to be useful for biological studies or as selection metrics in breeding programs (Figure 5). Heritability for manually measured phenotypes ranged from 0.26 (Stand Count) to 0.78 (Plant Height). PCA-based LSPs had higher heritability in the earlier PCs, which decreased rapidly to near zero after a dozen PCs. This pattern was expected because the earlier PCs capture greater amounts of variability than later PCs. The first sixteen PCs ranged in heritability from 0 (PC9 and PC12) to 0.44 (PC2). The sixteen LSPs produced by the autoencoder method do not have a logical ordering, the way that PCs do. In order to reference them they will be arbitrarily named ENC1 through ENC16. Autoencoder-based LSPs ranged in heritability from 0.04 (ENC2) to 0.41 (ENC1). The high-end heritabilities of the LSPs are comparable (±0.1) to many of the manually measured traits and high enough to be effectively selected on in a breeding program. These heritability measurements are from a relatively small sample (n=1,972) in a single environment, and are anticipated to rise as the rovers are deployed in replicated field trials.

**Figure 5:**
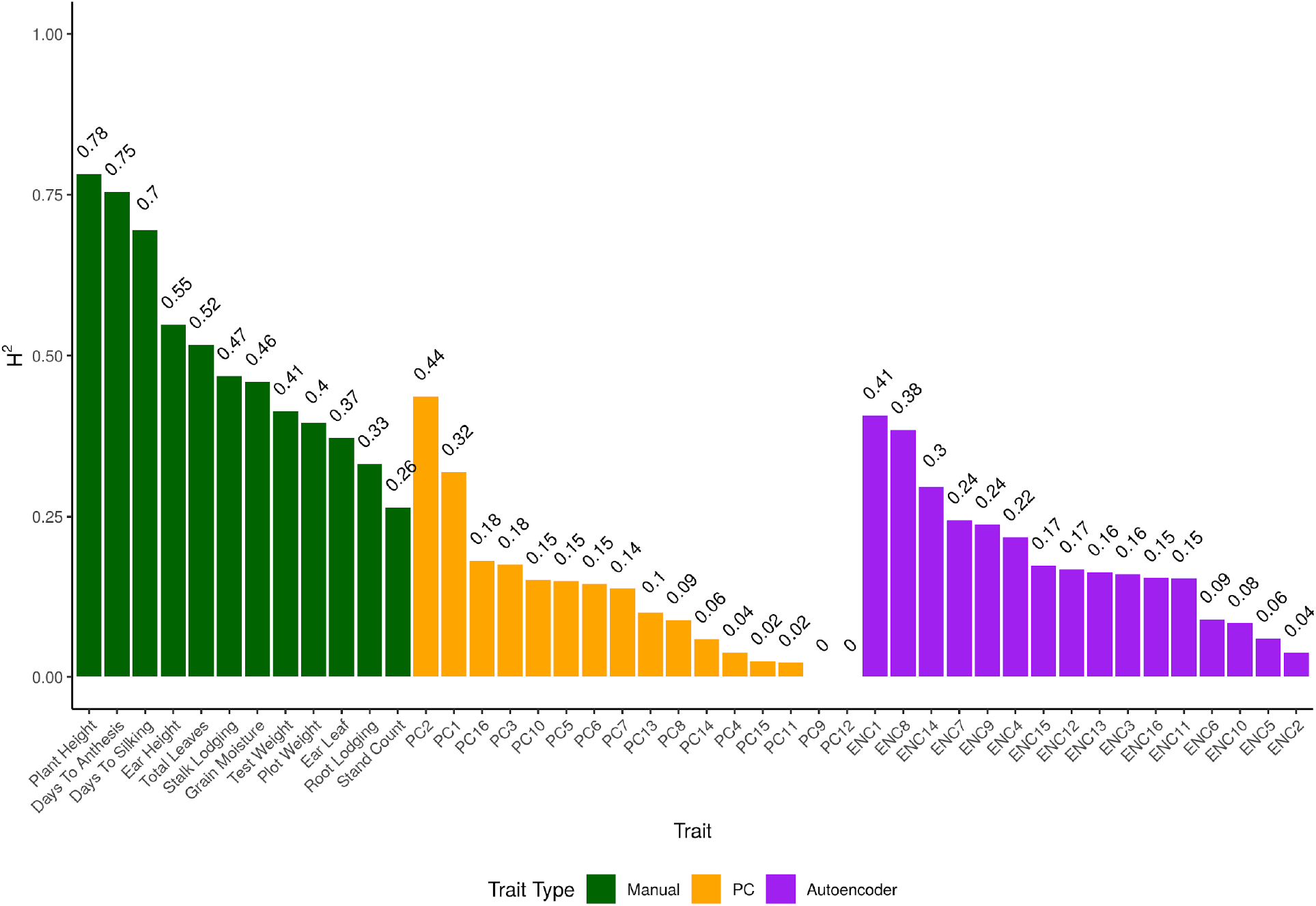
Comparison of heritability for manually measured traits and latent space phenotypes (LSPs). Manually measured traits (green) show a range of heritabilities. The higher-heritability LSPs (orange and green) have heritabilities similar to mid- and low-heritability manually measured traits. Heritability values are printed above the respective phenotype’s bar. All heritability measurements are from hybrid maize in a single environment.

### Predicting manually measured traits with Latent Space Phenotypes

Although the LSPs are heritable, that alone does not mean that they contain useful information about plant architecture. To determine whether the LSPs are capturing elements of biomass distribution and plant architecture, we used PLSR to predict each manually measured trait with either the first 16 PCA-based or all autoencoder-based LSPs (Figure 6). Prediction accuracy with PCA-based LSPs ranged from 0.25 (stalk lodging) to 0.89 (plant height), whereas prediction accuracy with autoencoder-based LSPs ranged from 0.24 (stalk lodging) to 0.85 (plant height). Similar prediction accuracies from autoencoder-based and PCA-based LSPs may be due to a limit in useful information contained in the plant surface density images. However, PCA-based LSPs are more predictive of manually measured traits if all PCA-based LSPs (rather than the first 16 PCs) are used for prediction. It is unsurprising that plant height showed the highest prediction accuracy, as it is straightforward to identify from the plant surface density images. Other architectural and agronomic traits, such as stand count, lodging, and leaf counts, demonstrated lower yet non-zero prediction accuracies. This could be for a few reasons: first, these traits do not show themselves as clearly as plant height in the image of plant surface distribution that was used as input for creating the LSPs; second, they are more difficult to measure manually and as such the measured values may be inaccurate due to human error during counting. Some traits, such as lodging, stand count, and all grain-related traits, were not yet fully determined when the rovers were driven through the field. For example, lodging and stand counts were done several weeks after the rover data were recorded, and may have changed in the intervening time. Generally, traits which were heritable and already stable or determined when rover data was collected (e.g., flowering traits, plant and ear height) had higher prediction accuracies from the LSPs than traits which had low heritability (e.g., leaf counts) or may have continued to change in the weeks after rover data were collected (e.g., stand count) (Supplemental Figure 1). As such, poor prediction accuracies may be due more to noisy manual measurements than to lack of information captured by LSPs.

**Figure 6:**
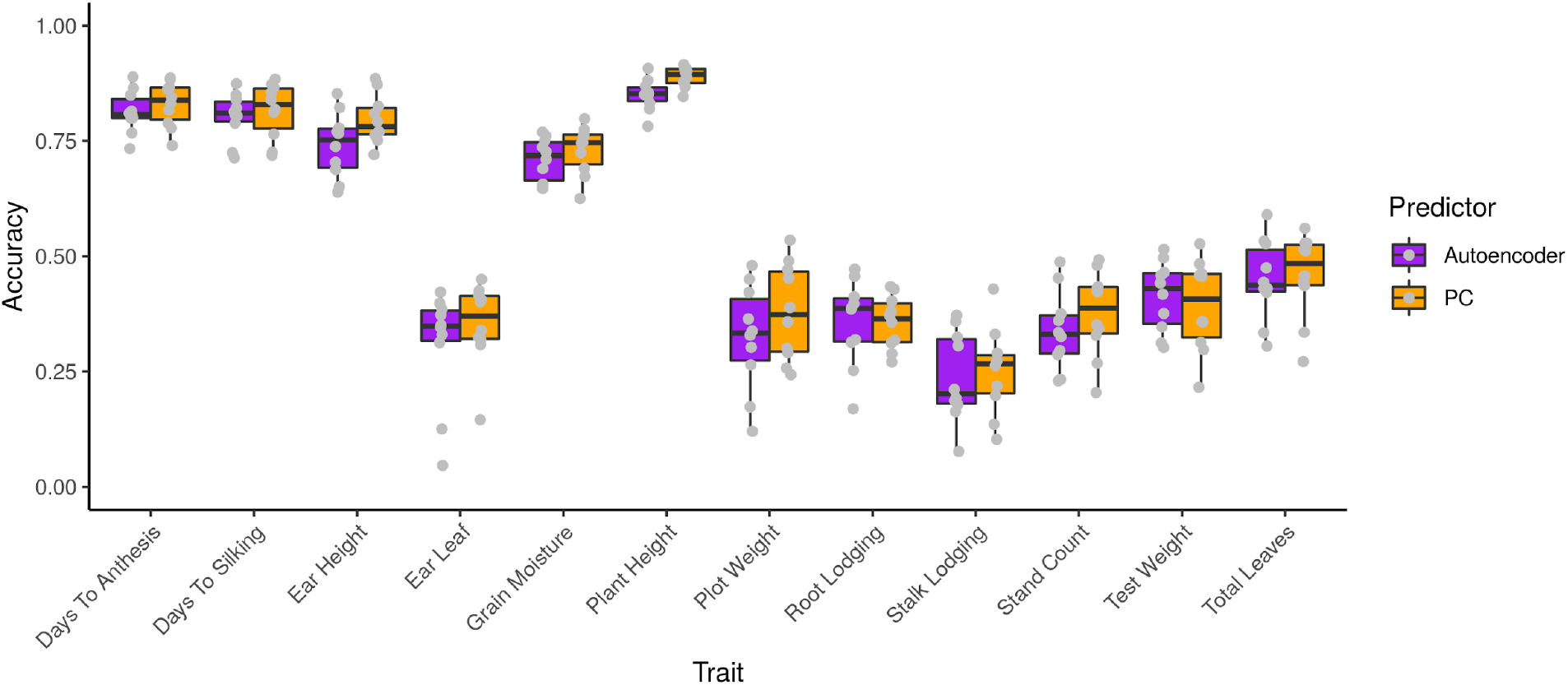
Ability of latent space phenotypes (LSPs) to predict manually measured traits. Autoencoder- and principal component analysis-based LSPs (purple and orange, respectively) were used to predict all manually measured traits by partial least squares regression. LSPs contain enough information about plant architecture to accurately predict architectural traits (Plant and ear height, ear leaf, total leaves, stalk and root lodging, and stand count), as well as non-architectural traits. Accuracy reported is the pearson correlation between predicted values and manually measured values.

LSPs were also predictive of non-architectural traits (grain moisture, test weight, plot weight, and flowering time) due to inherent correlations between them and agronomic or plant architectural traits. To demonstrate this, we used height, stand count, lodging, and leaf count traits as the explanatory variables in PLSR models to predict the non-architectural traits. The prediction accuracies of PLSR models trained on architectural traits are similar to the prediction accuracies of PLSR models trained on LSPs, showing that information about plant architecture is also indicative of non-architectural traits of agronomic importance (Supplemental Figure 2). Similarity in accuracy when predicting non-architectural traits with architectural traits versus LSPs shows that LSPs are capturing a similar quantity of information about plant architecture as the manually measured traits.

It should be noted that using LSPs to predict manually measured phenotypes is only meant to demonstrate that the LSPs contain signal related to plant architecture and performance. Higher prediction accuracy and better precision for such predictions can likely be achieved by creating algorithms or models that predict such traits directly from the LiDar data.

### Advantages, disadvantages, and future prospects for rover-based phenotyping

By showing that LSPs are both heritable and contain information about plant architecture, we have demonstrated that rover-based phenotyping and LSPs are promising for application to plant biology and crop breeding efforts. Sub-canopy rovers can be made to operate semi-autonomously, reducing the number of person-hours necessary to obtain quality phenotype data. The amount of time needed to manually gather phenotype data on a large, field-grown population is a function of the number of traits being recorded, the amount of time needed to measure and record each data point, the number of individuals helping with data collection, and the number of plots to be measured. The amount of time needed to measure a field by rover, on the other hand, is a function of the amount of time needed to drive through each plot and the number of plots to be measured. Because the LiDar enables three-dimensional reconstruction of each plot, all plant architectural information is recorded by default. The rover traverses the field at a rate of approximately 10 seconds per 5.33m plot. At that rate it can evaluate 2,000 two-row plots in less than 6 hours, easily achievable in a standard work day, and requires supervision by one person. If, instead, the same single person were to measure and record phenotypes manually on the same 2,000 plots, more time would be needed. Assuming an experienced researcher can collect phenotypic data at a rate of 20 seconds per plot, it would take between 11 and 12 hours to phenotype 2,000 plots. This is twice as long as it takes to evaluate the field by rover, and results in only a single datapoint per plot. If the researcher in this example is being paid $15 per hour, all of the rover traits presented here could be phenotyped for a cost of $0.04 per plot (for the researcher supervising the rover), whereas phenotyping a single trait manually would cost twice as much ($0.08) per plot. Considering the fact that the rover records enough data in a single pass to produce dozens of traits, whereas the cost of manually phenotyping scales linearly with the number of traits to be recorded, rover-based phenotyping quickly becomes orders of magnitude cheaper than manual phenotyping.

To rephrase this comparison, rover-based phenotyping allows calculation of a several LSPs with similar heritability to manually measured traits in the same amount of time needed to phenotype a single manually measured trait. If the purpose of phenotyping is to identify candidate genes via genetic mapping, rover-based phenotyping of LSPs could yield roughly a dozen times as many genetic mapping targets as manual phenotyping for the same investment of time. With the increasing throughput of gene editing technologies, screening such a large number of candidate genes may soon be a less daunting task than it is today (Ramstein et al., 2019). The relative ease of capturing data and processing it into LSPs can facilitate and standardize coordinated phenotyping efforts by research groups across a large number of locations. The same qualities can enable longitudinal evaluation of LSPs, opening the door to including developmental time as a standard axis of variation in experiments, similar to genotypic and environmental variation today.

Though rover-based phenotyping and LSPs show enormous promise for the fields of plant biology and breeding, they do still face some drawbacks. First and foremost, LSPs are of no benefit if they are not interpretable, useful, and applicable to biology or breeding. The results shown in this study provide evidence that LSPs meet these requirements, but more work is needed to prove their utility. The ultimate evidence will come with more data, when LSPs are used to identify candidate genes or as selection criteria in a breeding program. Although rover-based phenotyping has an immense advantage over manual phenotyping with regard to operating cost, it comes with the high up-front cost of purchasing the rovers themselves. In addition to startup cost, rover-based phenotyping is subject to the same pitfall as any other phenotyping system that records large quantities of information; methods need to be developed to analyze and interpret the data. LSPs are a convenient way to extract meaningful phenotypes from large quantities of complex and messy data. Scientists interested in faster or more accurate methods for measuring standard traits, such as the manually measured traits in this study, still need to develop reliable methods for doing so, which is a non-trivial task.

Though automated sub-canopy phenotyping comes with some challenges, the potential for benefit to plant biology and plant breeding communities is immense and real. The capabilities of sub-canopy rovers have been recognized by multiple groups (Mueller-Sim et al., 2017; Kayacan et al., 2018; Stager et al., 2019), and the promise of LiDar for in-field characterization of plant architecture is also recognized (Qiu et al., 2019; Su et al., 2019). By combining novel phenotyping and data analysis techniques, we have demonstrated that rover-based phenotyping by LiDar can produce LSPs that are heritable and contain information about plant architecture and plot-level biomass distribution. These techniques will enable in-field high-throughput phenotyping of crops in numerous locations and across developmental time by reducing the cost of collecting high-quality phenotypic data points.

### Data availability

The raw LiDar point clouds can be found at <DOI in preparation at CyVerse>. Code and phenotypic data are available at https://bitbucket.org/bucklerlab/p_lidar_lsp.

## Author Contributions

C.S, G.C., rover design and construction; J.L.G, E.S.B, M.A.G., study conceptualization; J.L.G, E.R., N.L, N.K, data collection; J.L.G., data analysis; all authors contributed to manuscript preparation or review.

## Acknowledgements

The information, data, or work presented herein was funded in part by the USDA-ARS, Genomes to Fields, and the Advanced Research Projects Agency-Energy (ARPA-E), U.S. Department of Energy, under Award Number DE-AR0000598. The views and opinions of authors expressed herein do not necessarily state or reflect those of the United States Government or any agency thereof. The use of trade, firm, or corporation names in this publication (or page) is for the information and convenience of the reader. Such use does not constitute an official endorsement or approval by the United States Department of Agriculture or the Agricultural Research Service of any product or service to the exclusion of others that may be suitable.

## Supplemental Material

**Supplemental Figure 1:**
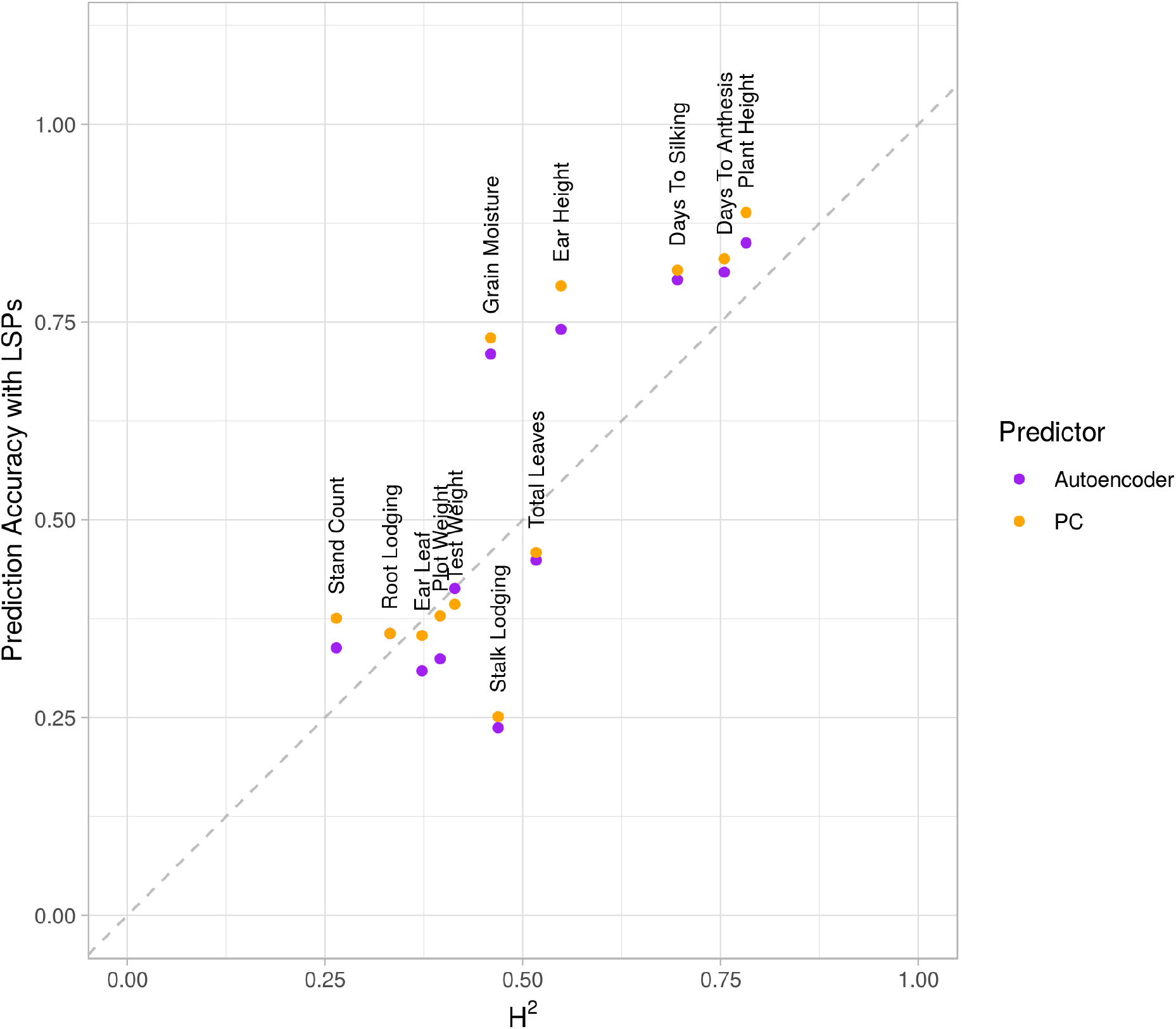
Correspondence between trait heritability (H^2^) and accuracy of predictions by latent space phenotypes (LSPs). Generally, LSPs are able to predict traits that are highly heritable, whereas LSPs have less predictive ability of traits with low heritability or that were measured weeks after rover data was collected (e.g., stand counts, lodging). Dashed line represents a 1:1 relationship between heritability and prediction accuracy.

**Supplemental Figure 2:**
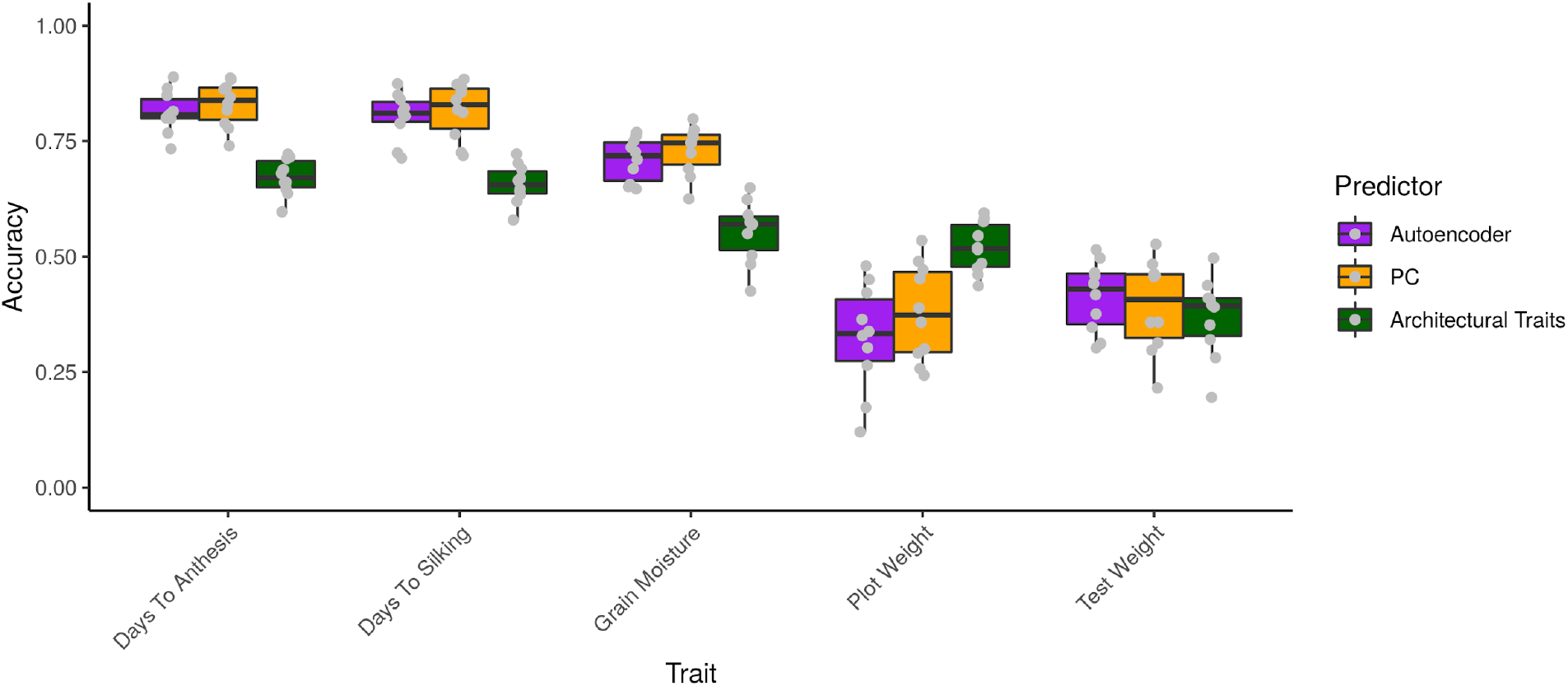
Prediction of non-architectural traits. Latent space phenotypes show high prediction accuracy of non-architectural traits, which are not directly visible to the LiDar. Architectural traits (plant and ear height, ear leaf, total leaves, stalk and root lodging, and stand count) were used to predict non-architectural traits by partial least squares regression, and compared to previously described results where LSPs were used as predictive variables. LSP results shown above (purple and orange) are identical to those in Figure 4. We found that architectural traits are predictive of non-architectural traits at a similar level of accuracy to LSPs.

